# Distinct Structural Features of the Lon Protease Drive Conserved Hand-over-Hand Substrate Translocation

**DOI:** 10.1101/617159

**Authors:** Mia Shin, Ananya Asmita, Cristina Puchades, Eric Adjei, R. Luke Wiseman, A. Wali Karzai, Gabriel C. Lander

## Abstract

Hand-over-hand translocation is emerging as the conserved mechanism by which ATP hydrolysis drives substrate translocation within the classical clade of AAA+ proteins. However, the operating principles of the distantly related HCLR clade, which includes the important quality control protease Lon, remains poorly defined. We determined a cryo-electron microscopy structure of *Y. pestis* Lon trapped in the act of processing substrate. This structure revealed that sequential ATP hydrolysis and hand-over-hand substrate translocation are conserved in this AAA+ protease. However, Lon processes substrates through a distinct molecular mechanism involving structural features unique to the HCLR clade. Our findings define a previously unobserved translocation mechanism that is likely conserved across HCLR proteins and reveal how fundamentally distinct structural configurations of distantly-related AAA+ enzymes can power hand-over-hand substrate translocation.

## Introduction

AAA+ enzymes (**A**TPases **a**ssociated with a variety of cellular **a**ctivities) constitute a broad superfamily of proteins defined by a structurally conserved domain that contains elements involved in nucleotide binding, sensing, and hydrolysis^1–4^. Despite structural similarities, AAA+ proteins play distinct roles in regulating diverse cellular activities including protein degradation, cytoskeleton remodeling, and DNA replication^1–4^. Importantly, AAA+ proteins have evolutionarily diverged into clades that are characterized by the incorporation of unique secondary structure elements into the canonical AAA+ domain^2,3^. Protein quality control serves as an excellent example of convergent functionality across distantly related clades of AAA+ proteins, as AAA+ proteases of both classical (i.e. FtsH and 26S proteasome) and HCLR (**H**slUV, **C**lpX, **L**on) families are required for maintaining protein homeostasis (or proteostasis) across all kingdoms of life^4^.

A fundamental question in the AAA+ field is whether distantly related proteins in distinct clades operate under a conserved mechanism to perform similar biological functions. Interestingly, recent cryo-electron microscopy (cryo-EM) structures of substrate-bound AAA+ proteins belonging to the classical clade revealed a unified mechanism for ATP-dependent substrate translocation^5–13^. In this mechanism, a spiral stair-case configuration of the ATPase domains gives rise to sequential, around-the-ring ATP hydrolysis and hand-over-hand substrate translocation. ATP-dependent quality control proteases in this clade display conservation of an inter-subunit signaling (ISS) motif that senses the nucleotide state of the neighboring subunit and transmits ATP-dependent conformational changes to substrate-interacting pore-loop residues within the central channel in the substrate translocation process^5–13^.

Interestingly, distinct structural components within the AAA+ domain that define the evolutionary divergent clades likely result in a diversification of the mechanisms utilized for function^2,3,14^. Consistent with this, AAA+ proteases belonging to the HCLR clade contain specific perturbations to key sequence motifs involved in the substrate translocation mechanism defined for classical AAA+ proteases. HCLR clade AAA+ proteases lack the ISS motif required for allosteric, nucleotide-dependent substrate translocation by classical proteases. Instead, HCLR proteases contain a pre-sensor-1 beta hairpin (PS1βH) insertion that is a defining element of the HCLR clade and has been shown to be critically important for substrate translocation^14^. While the PS1βH motif has been observed to directly contact DNA substrates in AAA+ helicases^15,16^, its position within protein translocases precludes substrate interaction, suggesting a different role for this motif in HCLR protease activity. Apart from the PS1βH, another notable distinction of the HCLR clade is the retention of the sensor-2 motif in helix 7 of the small ATPase subdomain, which is characteristically absent in classical AAA+ proteins^2^. These differences in sequence between HCLR and classical ATPases suggest distinct allosteric mechanisms of substrate translocation. However, the molecular implications of these differences in the activity of HCLR proteases remain unclear.

The highly conserved AAA+ protease Lon is a representative member of the HCLR clade that is responsible for maintaining proteostasis in diverse subcellular environments including the bacterial cytosol and the eukaryotic mitochondrial matrix^17–21^. Numerous biophysical approaches have been employed to gain insight into the specific nucleotide sensing and substrate translocation mechanism for this model HCLR protease^20–29^. Prior x-ray crystallography studies of ADP-bound *M. taiwanensis* Lon revealed a trimer-of-dimers configuration for the AAA+ domains wherein alternating sub-units were found to bind ADP^30^. This structure suggested a semi-concerted nucleotide hydrolysis mechanism involving coordinated movement of three alternating subunits that alternatively bind and hydrolyze ATP. However, this assertion is inconsistent with recent structural developments in the AAA+ field, notably the hand-over-hand substrate translocation mechanism observed for many classical AAA+ proteins.

Here, we present a cryo-EM structure of *Y. pestis* Lon trapped in the process of translocating substrate. Our Lon structure demonstrates that the hand-over-hand mode of substrate translocation is conserved in this HCLR protease. However, we show that Lon drives allosteric, nucleotide-dependent substrate translocation through a mechanism unique from previously described classical AAA+ proteins and dependent on the distinct structural features found in HCLR proteases. In this mechanism, the Lon PS1βH motif serves as an epicenter for allosteric regulation of substrate translocation, wherein nucleotide-dependent interactions with PS1βH coordinate rigid body movements of subdomains for around-the-ring ATP hydrolysis and substrate translocation. These results define a new molecular mechanism for hand-over-hand substrate translocation that is likely conserved throughout the HCLR clade. In addition, these findings shed light on the convergent evolution that occurred amongst diverse clades of AAA+ proteins to perform similar biological functions such as maintaining proteostasis through distinct molecular mechanisms.

## Results

### Structure of the substrate-bound *Y. pestis* Lon protease

For structural studies, we incubated full-length *Y. pestis* Lon bearing the slowly ATP hydrolyzing Walker B mutation (E424Q) with an excess of Y2853, an 18 kDa putative sensory transduction regulator protein that is a robust Lon substrate^31^. Substrate-bound complexes were separated from unbound substrate using size-exclusion chromatography (Figure S1A). Isolated complexes were incubated with saturating amounts of ATP (1 mM) and vitrified for single-particle cryo-EM analyses. This resulted in a reconstruction with an overall resolution of 3.4 Å (Figure S2A-E). The majority of the Lon N-domain (1-252) was not visualized in the reconstruction, likely due to conformational flexibility of this region. However, a 50 amino acid three-helix bundle (NTD^3H^) located directly N-terminal to the ATPase domains (253-305) was well-resolved (Figures 1A-C, 2A). The cryo-EM density of this bundle, as well as the ATPase motor and protease domains, was of sufficient quality for *de novo* atomic model building (Figure S2E-F). An additional density corresponding to an extended eight-residue polypeptide was identified in the central pore of the ATPase hexamer (Figures 1A, 2C). Although we were unable to confidently assign the identity of the amino acids or its polarity, we ascribe this additional density to a segment of the Y2853 substrate.

As expected, the secondary structural elements of the Lon protomer in our cryo-EM reconstruction resembles a previously determined crystal structure of substrate-free, ADP-bound *M. taiwanensis* Lon (PDB:4YPL) that included the NTD^3H^ (Figure S3). Individually, the NTD^3H^, ATPase, and protease domains of our ATP-bound subunit aligned well with a nucleotide-free subunit of *M. taiwanensis* Lon (0.78, 1.26, and 1.01 Å Cα RMSD, respectively). While conformational deviations are likely due to different loop conformations in the presence of substrate or distinct nucleotide states between the different protomers, these RMSD values emphasize conservation of secondary structural elements within these three domains between bacterial Lon homologs. However, our substrate-trapped structure offers a unique opportunity to dissect the molecular mechanisms of nucleotide sensing and substrate translocation utilized by the Lon protease.

### The AAA+ domains of substrate-bound Lon assemble into an asymmetric spiral staircase configuration

While the observed C6-symmetric organization of the serine protease domains in our cryo-EM structure is consistent with previously determined crystal structures, the quaternary organization of the ATPase domains is notably distinct. Whereas the crystal structure of *M. taiwanensis* Lon shows the NTD^3H^ and ATPase domains arranged in a symmetric “trimer-of-dimers” configuration, our reconstruction shows these domains assembled into an asymmetric spiral staircase with “seam” subunits positioned between the lowest and highest subunits of the staircase (Figure 1A-C). The ability of these ATPase domains to adopt such varied positions while covalently linked to a stable C6-symmetric protease is likely accommodated by a flexible 13 residue inter-domain linker containing a strictly conserved glycine residue located at the base of the small ATPase subdomain (G580, Figure S4A-B). This residue is analogous to the glycine linker between the ATPase and protease domains of other classical AAA+ mitochondrial ATPase proteases, previously shown to be critical for substrate translocation^5,32^. Consistent with a central role for this glycine residue in the mechanism of substrate translocation, incorporating a G580L mutation in Lon substantially diminished ATP hydrolysis and degradation of model substrates Y2853 and HspQ (Figure S4C-D).

The asymmetric configuration of the Lon subdomains spiraling around a centrally positioned polypeptide substrate is largely consistent with recently determined cryo-EM structures of other classical substrate-bound AAA+ protein translocases^5–13^. This structural similarity suggests that the staircase architecture is a conserved characteristic of the ATPase cassette in protein translocases across different clades and is indicative of a conserved substrate processing mechanism. Our cryo-EM structure provided the opportunity to explore how the unique structural motifs that distinguish the HCLR clade of AAA+ motors allosterically preserve, or depart from, the substrate processing mechanisms described for the classical clade.

**Figure 1.**
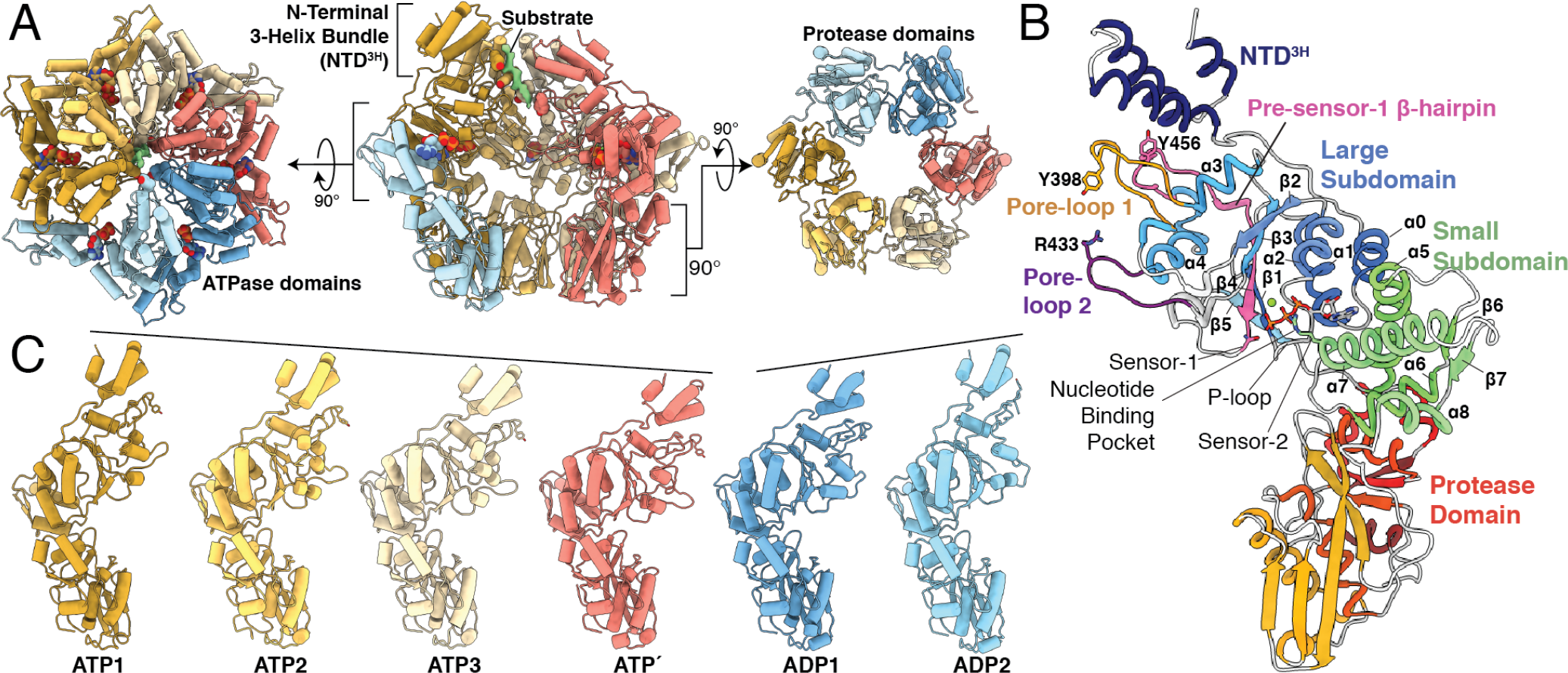
Architecture of the substrate-bound Lon protease. **A.** Cutaway view of the substrate-bound *Y. pestis* Lon atomic model (center) flanked by orthogonal views of the ATPase (left) and protease (right) domain rings. Cryo-EM density for substrate is colored green while each subunit of the homohexamer is assigned a color depending on its nucleotide state. ATP-bound subunits are colored different shades of yellow, the transitional ATP′ subunit is colored red, and ADP-bound subunits are colored in different shades of blue. Nucleotides and pore-loop 1 tyrosine residues are depicted using a sphere representation. **B.** A Lon protomer colored from N-to-C: the N-terminal 3-Helix bundle (NTD^3H^), large and small ATPase subdomains, and serine protease domain. Notable and conserved components of the Lon subunit are highlighted and/or labelled. Secondary structural elements of the AAA+ cassette are labelled using the canonical numbering for this domain^1^. **C.** Individual protomers oriented in the same direction, lined up side-by-side. Protomers were aligned using a secondary structural alignment of the protease domains. The descending and ascending movements of the NTD^3H^ and ATPase domains relative to their proteases are accentuated by solid lines shown above the NTD^3H^.

### Structure of Lon reveals a unique configuration of co-existing nucleotide states

A mixture of nucleotide states was previously proposed to establish the asymmetric organization of the ATPase ring utilized by AAA+ protein translocases to process substrates^5–13^. It is thought that structural rearrangements within the nucleotide pocket induced by ATP hydrolysis and exchange are allosterically propagated throughout the ATPase to coordinate substrate translocation. Similar to other AAA+ proteins^1–4^, the Lon nucleotide binding pocket is formed in the cleft between the large and small subdomains of the ATPase domain at the interface with the neighboring protomer (Figures 1B, S5). These pockets contain conserved structural motifs including the Walker A motif (356-GPPGVGKTS-364; important for nucleotide binding) and the Walker B motif (D423 and E424, mutated to Q in our structure); important for nucleotide hydrolysis)^1–4^. The Lon nucleotide binding pocket also contains a trans-acting arginine finger (R484), which is conserved in the classical clade of AAA+ enzymes, where it was shown to be critically involved in the sequential hydrolysis cycle^5–13^. Intriguingly, Lon, and indeed the entire HCLR clade of AAA+ proteins, additionally contains a cis-acting R542 sensor-2 residue that is absent from the classical clade^2^. Together with a sensor-1 residue, N473, these arginine residues are involved in positioning and stabilizing the ATP gamma phosphate for hydrolysis (Figures 1C, S5).

The cryo-EM density in the nucleotide binding pockets of our Lon reconstruction was of sufficient quality to unambiguously identify the nucleotide state in each of the six subunits (Figure S5). Four subunits are bound to ATP, and are hereafter named ATP1, ATP2, ATP3, and ATP′, in order from the uppermost subunit of the ATPase staircase to the lowest (Figure 1C). Our structure further revealed two ADP-bound “seam” subunits (hereafter named ADP1 and ADP2) that are somewhat displaced from the hexamer, with ADP1 at an intermediate position between the lowermost and uppermost staircase subunits, and ADP2 at approximately the same height as ATP1 (Figure 1C). The presence of two ‘seam’ subunits observed in the Lon hexamer is in contrast to cryo-EM structures of classical AAA+ proteins that show five closely-associated staircase subunits with only one “seam” subunit displaced from the spiral staircase organization. Thus, the unique configuration of Lon subunits suggests that this AAA+ protease could show distinct differences in substrate engagement, as compared to classical AAA+ proteases.

### Nucleotide state does not directly dictate pore-loop interaction with substrate

Recent cryo-EM structures of substrate-bound AAA+ proteins have shown that pore-loop aromatic residues protrude from ATP-bound subunits into the central channel of the spiral staircase to directly interact with substrates via sequence-independent backbone intercalations^5–13^. Structures of classical clade AAA+ proteins have established a direct correlation between nucleotide state, pore-loop conformation, and inter-actions with substrate. Whereas all ATP-bound subunits in prior structures of AAA+ protein translocases display nearly identical pore-loop arrangements and substrate interactions, our Lon structure shows that substrate interactions between the aromatic pore-loop 1 residue (Y398), previously shown to be important for substrate translocation^5–13^, does not directly correlate with subunit nucleotide state. The Y398 residues of the topmost three descending ATP bound subunits in the spiral staircase form tight interactions with the unfolded substrate backbone (Figure 2A, C-D). Intriguingly, despite containing ATP in the nucleotide-binding pocket, the Y398 pore-loop 1 residue in the lowest subunit of the spiral, ATP′, is displaced from the central pore and does not contact substrate (Figure 2C). This architectural disparity with all other substrate-bound AAA+ protein translocase structures is also observed in the ADP-bound subunits. Whereas the pore-loop 1 aromatic of the lowermost ADP-bound subunit from classical AAA+ proteases has previously been observed in proximity to substrate^5–13^, the pore-loops of both ADP-bound seam sub-units in Lon are completely disengaged from substrate (Figure 2C). This observation that Lon engages substrates through direct interactions with only three pore-loop 1 aromatic residues, despite containing four ATP-bound subunits, indicates that these interactions do not directly correlate with nucleotide binding state. Thus, this likely represents an allosteric departure from the mechanism proposed for classical AAA+ proteases.

**Figure 2.**
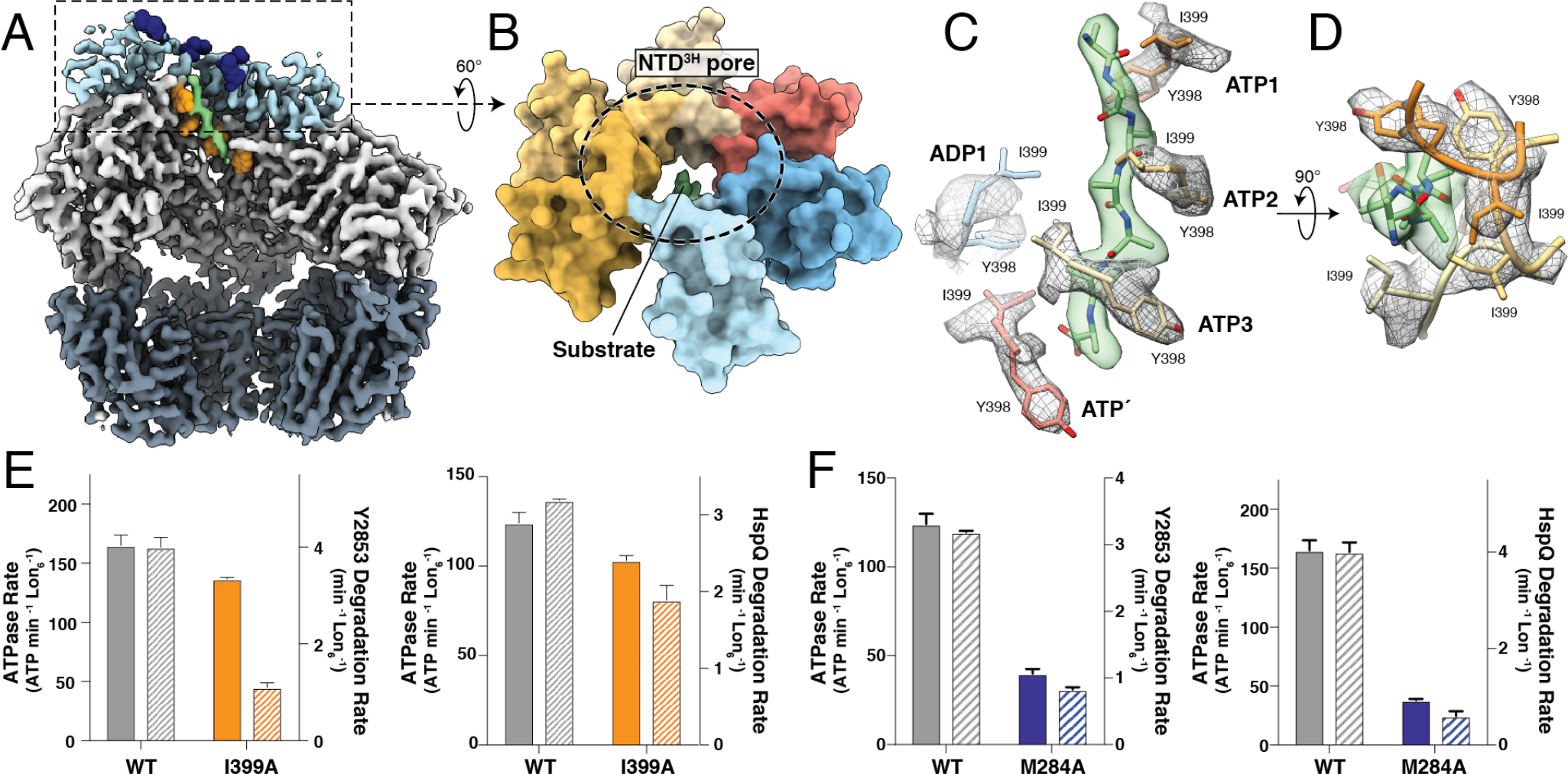
Substrate translocation is mediated by residues in pore-loop 1, pore-loop 2, and the NTD^3H^. **A.** Cutaway view of the substrate-bound Lon cryo-EM density with NTD^3H^ colored light blue, ATPase domains colored light grey, protease domains colored dark grey, and substrate colored green. Sphere representations of spiraling methionine (M284) side chains in the NTD^3H^ and tyrosine residues of pore-loop 1 are colored dark-blue and orange, respectively. **B.** The NTD^3H^ pore, shown as a molecular surface representation and colored by subunit as in Figure 1A. **C.** An eight-residue polyalanine chain is modeled into the substrate density, shown in a transparent green surface representation. Y398 and I399 from pore-loop 1 are shown using stick representations with associated cryo-EM density zoned around these residues in gray. While Y398 and I399 show intercalating, zipper-like interactions with substrate in ATP1-3, Y398 and I399 of pore-loop 1 in ATP′ are positioned further away from the central channel. Pore-loop 1 residues in the seam subunits (ADP1 and ADP2) are completely removed from substrate. **D.** An orthogonal view of Y398 and I399 interactions with substrate emphasizing “pincer-like” gripping of the substrate by these two hydrophobic residues. **E.** Mutating pincer residue I399 in the conserved pore-loop 1 marginally affects ATP hydrolysis, but decreases degradation of known Lon substrates, Y2853 and HspQ. **F.** Lon bearing a M284A mutation in the NTD^3H^ shows a defect in both ATPase and degradation rates for both Y2853 and HspQ substrates.

Intriguingly, our structure shows that the hydrophobic pore-loop 1 residue I399, which neighbors the aromatic Y398, contributes an additional intercalating interaction with substrate. Together, these two pore-loop 1 residues in Lon flank both sides of the incoming substrate, engaging the polypeptide backbone with a pincer-like grasp to facilitate translocation into the peptidase chamber (Figure 2C-D). Consistent with a key role in substrate translocation, the I399A mutant of Lon exhibits a severe defect in the degradation of the Lon substrates HspQ and Y2853, while only minimally impacting ATP hydrolysis (Figure 2E). This suggests that this double-intercalation mechanism may have evolved to strengthen interaction with substrate, given that only half of the pore-loops in the hexamer appear to engage substrate in bacterial Lon species.

### Spiraling N-terminal helical bundles play a critical role in substrate processing

Whereas a spiraling organization of pore-loops appears to be the generally conserved manner in which AAA+ enzymes interact with protein substrates, recruitment and transfer of substrates to the ATPase domains is primarily accomplished through an evolutionarily diverse range of structural components. In Lon, substrate recruitment is likely accomplished by the 300-residue N-terminus, which contains an N-terminal beta-sheet domain connected to a long alpha helix, followed by the NTD^3H^ ^33,34^. The long alpha helices of neighboring subunits are thought to assemble into coiled-coils, resulting in a trimer-of-dimer arrangement of the N-terminal regions in the full-length structure^35^. Truncating the entire N-terminal domain in several Lon homologs ablates both ATPase and proteolytic activity, but these activities can be restored by inclusion of the NTD^3H^ ^36^. However, substrate-stimulated ATPase and protease rates are diminished in Lon that only includes the NTD^3H^ of the N-terminus^36,37^, suggesting that this domain may function as an intermediary between the regions of Lon that are involved in substrate recruitment and translocation.

The NTD^3H^s of Lon were previously proposed to be flexibly attached to the ATPase domain, given that these domains are connected by a 12-residue linker^27^. However, our structure shows that the NTD^3H^s are rigidly positioned above the ATPase domains and tightly incorporated into the staircase organization, forming a pore approximately 15 Å in diameter (Figure 2B). The extensive interactions between the ATPase domain and the NTD^3H^ suggest that the NTD^3H^ likely functions as a rigid extension of the ATPase domain. We thus speculated that the NTD^3H^ might be directly involved in substrate translocation.

While we do not observe substrate density extending into the NTD^3H^ staircase (Figure 2A-B), our structure demonstrates that the ATP-dependent spiraling organization of the ATPase domains can be transmitted to regulatory domains attached to the enzyme. Our Lon reconstruction showed that residue M284 is positioned at the inter-domain interface within a helix-turn at the entrance to the NTD^3H^ pore. This uniquely positions this residue such that it could simultaneously be involved in substrate interactions as well as stabilizing the pore-like organization of the NTD^3H^s (Figure 2B). We tested the relevance of this methionine in substrate processing by introducing an M284A mutation. We confirmed that the mutant is fully competent in forming a substrate-bound hexamer by size-exclusion chromatography (Figure S1B). However, this mutation significantly reduced substrate-stimulated ATPase and protease activity, implicating M284 in the LON substrate translocation mechanism (Figure 2F). Given its location at the entrance to the NTD^3H^ pore (blue in Figure 2A), these data suggest that M284 is involved in guiding targeted polypeptides from the N-terminal substrate-recruiting regions of the Lon N-terminus to the pore-loops within the central ATPase channel for unfolding and translocation, either through direct substrate interactions or through stabilization of the NTD^3H^ pore.

### Hand-over-hand substrate translocation is conserved in the HCLR protease Lon

The hand-over-hand mechanism of AAA+ substrate translocation is defined by a staircase of conserved aromatic pore-loop residues arranged in a spiral staircase within the central ATPase channel, whose interactions with substrate are allosterically influenced by the nucleotide state as sequential, around-the-ring ATP hydrolysis occurs within the hexamer. We have established that the overall spiraling organization of our Lon structure is generally consistent with other cryo-EM structures of substrate-bound classical AAA+ protein translocases, thus supporting a conserved hand-over-hand substrate translocation mechanism among AAA+ protein translocases across diverse clades. However, all classical AAA+ proteases contain a dynamic trans-acting ISS-motif positioned near the nucleotide binding pocket at the inter-subunit interface^5,7^. This motif alternates between an extended conformation and an alpha helical fold in a nucleotide-dependent manner to dictate pore-loop interactions with the translocating substrate. As such, the ISS plays a critical role in establishing the substrate translocation mechanism for classical AAA+ protein translocases. However, the ISS motif is not present in Lon or any other HCLR clade AAA+ protease^2,14^. Our cryo-EM structure of Lon, which reveals a previously unobserved structural arrangement of ATPase domains and pore-loops, provides the opportunity to describe precisely how the unique structural motifs that distinguish the HCLR clade of AAA+ motors are allosterically regulated to drive this conserved hand-over-hand mechanism.

### The PS1βH connects nucleotide-sensing elements to the substrate-interacting regions

The PS1βH motif is the defining feature of the HCLR clade of AAA+ proteins^2,3,7,14^. Thus, we sought to define the specific involvement of this motif in Lon-dependent hand-over-hand substrate translocation. The PS1βH runs laterally along the inter-subunit interface of Lon, with the turn of the hairpin positioned in close proximity to pore-loop 1 (Figure 3A-D). A conserved tyrosine residue (Y456) within the PS1βH turn inserts into a pocket at the interface of the NTD^3H^ and pore-loop 1 and is stabilized by conserved tryptophan (W297) and tyrosine (Y294) residues (Figure 3B). Additionally, a conserved glutamate residue (E458) from PS1βH interacts with two tandem arginine residues (R395 and R396) that are immediately adjacent to the conserved pore-loop 1 aromatic residue (Y398) (Figure 3C). This suggests an important coordinating role for the PS1βH and the position of the substrate-interacting pore-loop 1 within the protomer. The importance of this structural coordination was confirmed by mutagenesis. An E458A mutation impaired degradation of the Lon substrates Y2853 and HspQ but did not substantially impact ATPase activity (Figure 3E). This supports a model whereby the charged interactions between R395/396 and E458 of pore-loop 1 and the PS1βH stabilize pore-loop 1 interactions with substrate.

**Figure 3.**
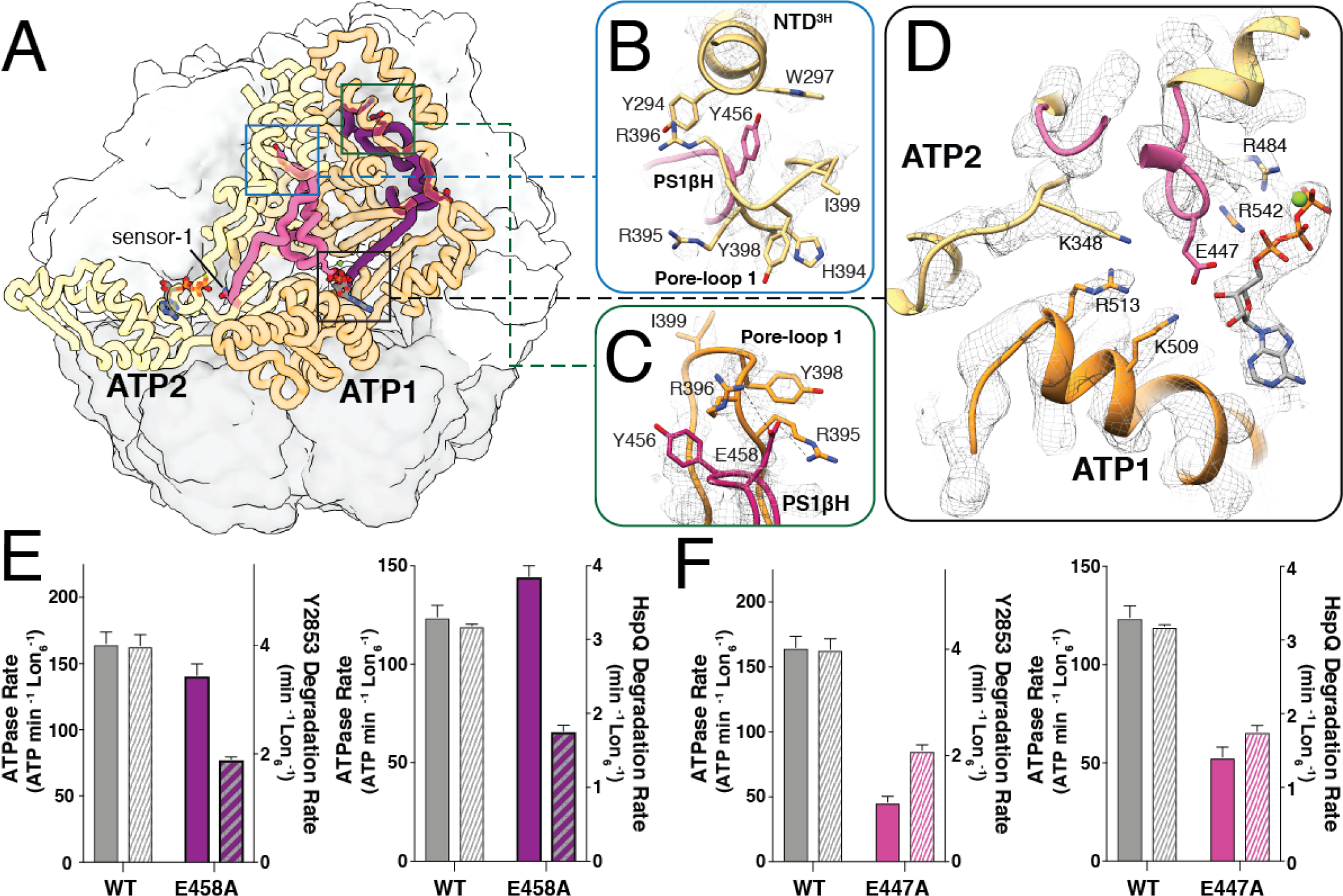
The PS1βH motif of Lon connects pore loop 1 to adjacent nucleotide binding pockets. **A.** ATP1 and ATP2 subunits are highlighted using a coil representation in the context of Lon structures (smoothed surface representation, ATPase domains white and protease domains gray). The PS1βHs are colored purple and pink in ATP1 and ATP2, respectively. Sensor-1 at the C-terminal base of the PS1βH is denoted in ATP2. Close-ups of the regions enclosed by the boxes are shown in **(B-D)**. **B.** Y456 at turn of the PS1βH stabilizes the NTD^3H^ through proximal interactions with conserved aromatic residues Y294 and W297. **C.** Pore-loop 1 interactions with substrate are stabilized through interactions between E458 in the PS1βH and two tandem arginine residues in pore-loop 1. **D.** The N-terminal base of PS1βH interacts with the nucleotide binding pocket in *trans* using a bridging glutamate (E447) that is stabilized by a patch of basic residues in the counterclockwise adjacent subunit. **E.** Mutating E458 to alanine shows a minor effect on ATPase activity but a significant reduction in degradation of known Lon substrates, Y2853 and HspQ, suggesting that E458 is indirectly involved in pore-loop 1 interaction with substrate. **F.** Mutating the glutamate bridge E447 to alanine substantially diminishes both ATPase activity and substrate degradation.

In addition to stabilizing pore-loop 1 within the protomer, the PS1βH is structured to sense or influence the nucleotide state within the subunit as well as that of the preceding protomer in the staircase. The C-terminal end of the PS1βH continues into the β4 strand of the canonical AAA+ fold that contains the sensor-1 residue (N473) involved in ATP hydrolysis within the protomer (Figures 1B, 3A, S5–6). This organization of the PS1βH is thus sensitive to the nucleotide states at either end of this motif and, given the role that the PS1βH plays in stabilizing pore-loop 1, likely plays a central role in allosterically coordinating substrate translocation. Interestingly, at the other end of the PS1βH, a highly conserved glutamate residue (E447) extends towards the nucleotide binding pocket of the neighboring subunit. Within the uppermost three ATP-bound binding pockets, this glutamate interacts with a patch of basic residues within the adjacent small subdomain that includes the sensor-2 residue R542 (Figure 3D). However, within the other three nucleotide binding pockets, this transacting glutamate is retracted from this basic patch (Figure 5B). We speculated that this charged interaction is important for inter-subunit stabilization, and that its disruption may be result in displacement of protomers from the hexamer. We investigated the relevance of this “bridging glutamate” residue in the Lon mechanochemical cycle by introducing an E447A mutation, which caused severe defects in both substrate-induced ATPase and proteolytic activity (Figure 3F).

Together, these interactions tightly link the PS1βH element to both the nucleotide binding pocket and the critical substrate-interacting regions in Lon. These findings demonstrate that the conserved PS1βH functions as a structural entity linking cis- and trans-acting nucleotide state sensors to the substrate translocation elements. Notably, an alignment of the PS1βH from crystal structures of HslUV, ClpX, and RuvB and their respective sequences shows a conserved position of this glutamate residue at the N-terminal base of the hairpin (Figure S7A-B). The structural and sequence conservation of this bridging glutamate residue suggests that its mechanistic importance is likely conserved throughout the HCLR clade of AAA+ proteins. Given the relevance of these structural elements to Lon function, we next examined how the PS1βH motif structurally transitions through the hydrolysis cycle.

**Figure 4.**
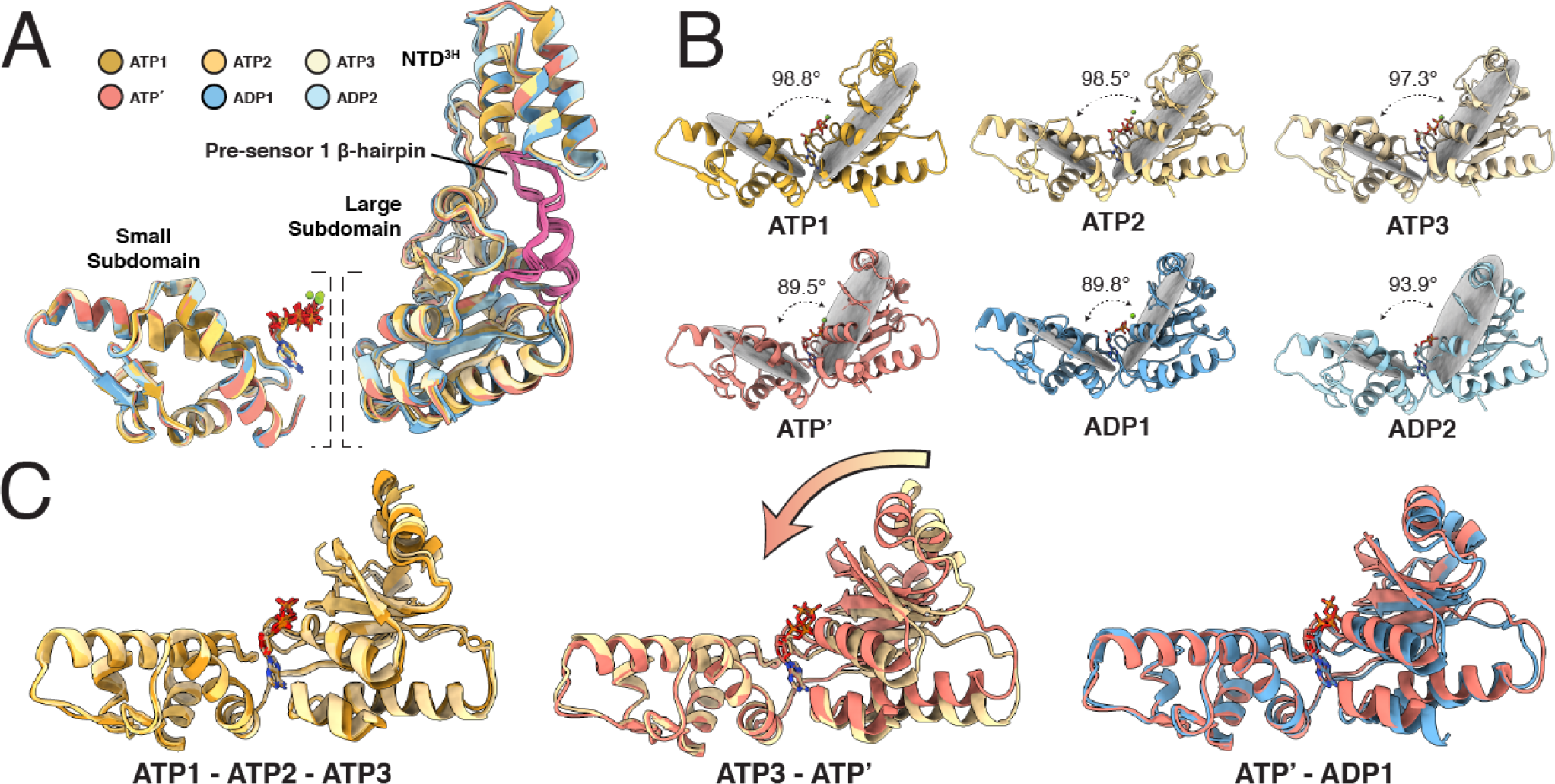
Rigid body rearrangements of the ATPase domains. **A.** Secondary structure-based alignment of individual units of the AAA+ domains, including the NTD^3H^ and large and small ATPase subdomains. Each subunit is colored according to the same color scheme assigned in Figure 1. Superimposing subdomains reveals an average CαRMSD of 0.81, indicating that these regions move as rigid units throughout the hydrolysis cycle. **B.** Individual ATPase subdomains oriented in the same direction, lined up side-by-side. ATPases were aligned based on a secondary structural centered on the small ATPase subdomain. Dihedral angles between the large and small ATPase subdomains are shown using gray planes, showing compression of the ATPase domains in the ATP′ and ADP1 subunits. **C.** Secondary structure alignment of ATPase subunits based on the small ATPase subdomains. While the three topmost subunits of the ATPase spiral staircase are in similar configurations (**left**), the ATP′ subunit is in a compressed state (**center**) that is retained upon ATP hydrolysis and phosphate release in the ADP1 subunit (**right**).

### Rigid body motions of the ATPase subdomains drive substrate translocation

It is widely accepted that an around-the-ring mechanism of substrate translocation within classical AAA+ proteases requires that loss of a gamma phosphate induced by nucleotide hydrolysis results in local structural rearrangements that are propagated to neighboring subunits^5–13^. However, in contrast to the ISS motif found in classical AAA+ proteases, our structure shows that the nucleotide sensing elements of the PS1βH described above do not undergo nucleotide-dependent rearrangements of local secondary structure at the inter-subunit interface. In fact, superimposing the ATPase large subdomain of the six Lon subunits (independent of the small subdomain) reveals an average Cα RMSD of 0.81 Å (Figure 4A). Similarly, independently superimposing the small ATPase subdomains also showed that these regions were structurally consistent within the six protomers of the Lon hexamer (Cα RMSD of 0.44 Å) (Figure 4A). Regardless, since the PS1βH appears to stabilize pore-loop 1 while not directly contacting substrate, it is likely that the PS1βH plays an important stabilizing role in coordinating hand-over-hand translocation as the subunits move as rigid bodies through a previously undescribed substrate translocation mechanism.

Interestingly, and consistent with all prior cryo-EM structures of substrate-bound AAA+ translocases, the dihedral angles between the large and small ATPase subdomains fluctuate within the hexamer (Figure 4B). Expectedly, given the shared nucleotide state, the dihedrals between the large and small ATPase subdomains in the ATP-bound subunits ATP1, ATP2, and ATP3 are indistinguishable in our structure. However, the fourth ATP-bound subunit, ATP′, displays a conspicuous 9° closure of the subdomains (Figure 4A-B). This distinct conformational arrangement of the lower-most ATP-bound subunit of the ATPase staircase indicates that the organization of the Lon nucleotide-binding pocket can accommodate two different ATP-bound states. Moreover, the dihedrals between the small and large subdomains of the ATP′ and ADP1 are almost unchanged, giving rise to a nearly identical organization of the ATPase domain despite containing different nucleotide states (Figure 4B).

To help identify the mechanism of allostery responsible for Lon substrate processing, we linearly interpolated between the subunits of the homohexamer as a means of approximating an around-the-ring ATP hydrolysis cycle and observed the motions of the large and small ATPase subdomains as they progress through the mechanochemical cycle (Figure 4C, Movie 1). Intriguingly, the ADP2 subunit remains in a nearly fixed position within the hexamer as ADP is exchanged for ATP within the protomer, showing no noticeable movement relative to protease domains (Figure 1B, Movie 1). Rather, nucleotide exchange is concurrent with a concerted motion of the ATP-bound subunits, which pivot as a rigid body approximately 12° towards the central channel (Figure 4C, right) to form inter-subunit interactions with the newly formed ATP1 subunit. Exchange of ADP for ATP within a subunit causes the arginine finger and bridging glutamate E447 of the adjacent subunit to stabilize inter-subunit interactions with the ATP phosphates and a patch of basic residues (K348 and R484 in *cis* and R513, K509, R542 in *trans*) within the newly ATP-bound subunit, respectively. Notably, this rocking of the ATP-bound subunits results in the positioning of the substrate for interaction with the pincer-like pore-loop 1 residues of the newly formed ATP1 subunit at the top of the spiral staircase, while simultaneously translocating the bound substrate the length of two amino acids toward the protease domains.

Notably, the pivoting movement of the ATP-bound subunits coincides with a collapse of the large and small sub-domains within the lowermost ATP-bound protomer of the spiral staircase (Fig 4B, Movie 1). This motion of the large subdomain towards the small subdomain would result in a substantial steric clash with the two ADP-bound subunits (Movie 1). As a result, these subunits become displaced from the ATPase hexamer, forming the two ADP-bound “seam” subunits (Movie 1).

### Structure-based mechanism for Lon-dependent hand-over-hand substrate translocation

In Lon, ATP binding causes rigid body motions of the ATPase subdomains to confer a hand-over-hand mode of substrate translocation mediated by the PS1βH insertion (Figure 5A). In the ATP1, ATP2, and ATP3 subunits, the ATP nucleotide is stabilized through interactions of the trans-acting arginine finger (R484) and cis-acting sensor-2 (R542) with the β-γ phosphates of ATP (Figures 3D, 5B, S5). The trans glutamate (E447) at the N-terminal base of the PS1βH, that is important for the ATP hydrolysis cycle, extends towards the nucleotide binding pocket, engaging with a cluster of basic residues (Figures 3D, 5B). Curiously, the trans-acting nucleotide-stabilizing interactions present in the three ATP-bound subunits are abolished in the ATP′ subunit, despite the presence of a bound ATP nucleotide (Figures 5, S5). In this subunit, the β-γ phosphates of ATP are primarily stabilized by the cis-acting sensor 2 arginine that is not present in classical AAA+ proteins (Figures 3D, S5).

The loss of inter-subunit contacts is likely due to ATP binding and steric clashes occurring between the four ATP-bound subunits, which results in a collapse of the ATP′ nucleotide binding pocket and displacement of the adjacent ADP-bound ATPase domain (ADP1) from the spiral staircase (Fig 4A-B, Movie 1). As the trans-acting arginine finger retracts from the ATP′ binding pocket, sensor-2 arginine draws closer to the β-γ phosphates of bound ATP (Figure 5B-C). Closure of the binding pocket simultaneously draws the sensor-1 and Walker B elements toward the ATP nucleotide, which likely “primes” this subunit for the next ATP hydrolysis event.

Atop the ATPase domains, the NTD^3H^s exhibit a striking gyration as a result of these rigid body motions, giving rise to a counterclockwise movement of the NTD^3H^ pore. Notably, the four NTD^3H^s of the ATP-bound subunits are in close proximity to one another and move as a nearly rigid body. This is in accordance with our prior observation that the NTD^3H^ is tightly associated with the ATPase large subdomain and functions as a structural extension of this region. It is likely that the rigid body motions associated with nucleotide exchange and ATP hydrolysis are amplified through the NTD^3H^ and further promote displacement of the ADP-bound subunits to accommodate the collapse of the ATP′ subunit and ATP hydrolysis. This observation that the NTD^3H^ may be involved in repositioning the ATPase domains during the hydrolysis cycle would explain why constructs lacking the NTD^3H^ show a complete loss of enzymatic function.

## Discussion

Our reconstruction of the Lon protease reveals that the hand-over-hand model for substrate translocation is conserved in this AAA+ protein of the HCLR clade. Moreover, we define a mechanism for nucleotide-dependent substrate translocation distinct from that currently proposed for the AAA+ proteases belonging to the classical clade of ATPases. Rather than involving nucleotide hydrolysis-induced rearrangements of local secondary structure that are allosterically communicated to substrate-interacting pore loops, Lon appears to process substrates through rigid-body motions that are mediated by structural features that define the HCLR clade of AAA+ proteins: the PS1βH and a sensor-2 arginine. The PS1βH spans the ATPase large domain and plays a critical role in ATP binding and hydrolysis via sensor 1, located at the N-terminal end of the PS1βH motif. The PS1βH also establishes inter-subunit interactions through a bridging glutamate located at the N-terminal end of the hairpin. The turn of the hairpin, located sequentially and spatially between these elements, stabilizes the substrate-interacting pore loops at the center of the ATPase channel. The sensor-2 arginine within the small ATPase subdomain likely plays an important role in coordinating ATP for hydrolysis within the lowermost subunit of the staircase.

Numerous structural rearrangements must occur simultaneously or in close succession in order to accommodate the substrate translocation mechanism in Lon. ATP binding establishes inter-subunit interactions that result in a pivoting motion of one half of the ATPase hexamer, positioning pore loop residues for substrate interaction at the top of the staircase while translocating the unfolded polypeptide toward the proteolytic chamber. These motions are accommodated by a displacement of the ADP-bound subunits from the hexamer, which disrupts the inter-subunit interactions between the lowermost subunits of the staircase. Perturbation of these stabilizing interactions coincides with a compression of the lowest nucleotide binding pocket, priming the bound ATP for hydrolysis. These concerted motions are coordinated by the PS1βH, which connects the bridging glutamate, pore loops, and sensor-1.

**Figure 5.**
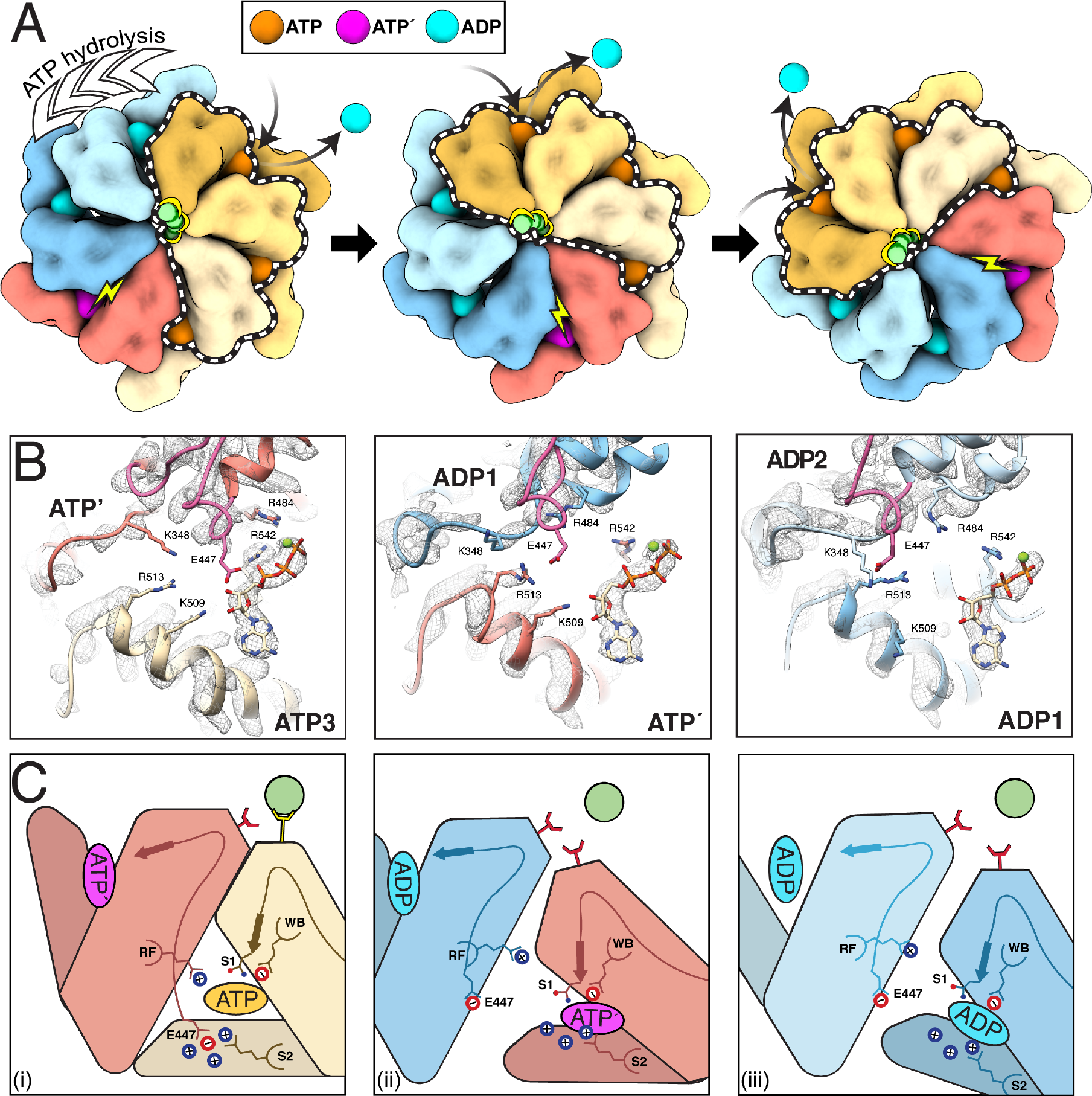
Summary of the allosteric mechanism of substrate translocation in Lon protease. **A.** Around-the-ring, counter-clockwise ATP hydrolysis drives interactions between substrate (green) and pore loop residues within three ATP-bound AAA+ domains. Nucleotide exchange in the second ADP-bound ‘seam’ subunit approximated by a 60° rotation of our structure results in lateral movements of the three upper-most ATP bound subunits (dotted outline). These lateral movements lead to a two amino acid step in substrate translocation and ‘primes’ the now ATP′ subunit (red) for ATP hydrolysis (see also Movie 1). **B.** Remodeling cis and trans subunit interactions in the nucleotide binding pocket drives sequential ATP hydrolysis cycle and stepwise substrate translocation. In the nucleotide binding pocket of two adjacent ATP-bound subunits (e.g, ATP1-ATP2, left), nucleotide is stabilized by interactions with the arginine finger (R484) in *trans* and sensor 2 (R542) in *cis*. Furthermore, a bridging glutamate residue (E447) at the N-terminal base of the PS1βH motif engages a cluster of basic residues within the nucleotide-binding pocket, stabilizing the inter-subunit interface. In the ATP′ nucleotide binding pocket (middle), this organization is disrupted upon E447 being retracted from the nucleotide binding pocket, causing the subunit to compress, bringing sensor-2 and other motifs involved in ATP hydrolysis (i.e. Walker A and Walker B motifs) in the proximity of the bound nucleotide. The ADP1 (right) and ADP2 (not shown) nucleotide binding pockets reveal a similar organization to that observed in ATP′, indicating that nucleotide exchange is necessary to ‘reset’ the hydrolysis cycle. **C.** Cartoon summary of the mechanism of allostery involved in hand-over-hand substrate translocation in Lon, consisting of three parts: (i) Substrate interactions in three ATP-bound subunits dictated by a trans-acting ‘bridging glutamate’ in PS1βH that is stabilized by an acidic patch in the nucleotide binding pocket. These inter-subunit charged interactions determine pore-loop 1 interactions with substrate. ADP release and ATP binding at the top of the spiral staircase causes (ii) rigid body rotations of ATP-bound subunits down the spiral staircase, leading the ATP′ subunit to compress, be “primed” for the next ATP hydrolysis event, and detach from bound substrate. Simultaneously, (iii) the ADP1 subunit is released from the spiral while maintaining a compressed ATPase configuration. See also Movie 1.

We have demonstrated that the unique elements that characterize the HCLR clade of AAA+ proteins are critical for the substrate processing mechanism of Lon. We thus posit that the around-the-ring hydrolysis mechanism defined here is likely conserved throughout the HCLR clade, including ClpXP and HslUV enzymes. Moreover, the hydrophobic-aromatic pore loop 1 conserved in AAA+ proteases of the HCLR is absent in RuvB, a DNA translocase and HCLR clade member. It is likely that RuvB has evolved divergent functionalities of the PS1βH to interact directly with DNA substrates meanwhile utilizing the same allosteric mechanism as Lon centralized around this conserved motif to transmit conformational changes across the hexamer.

Together, these findings establish a structural basis to explain the convergent mechanisms by which evolutionarily distinct AAA+ proteases can perform similar biologic functions such as proteostasis maintenance. AAA+ proteins in different clades of AAA+ arose early on in evolution, and each developed unique structural features to accomplish a myriad of biological functions with diverse target substrates^2,3^. In certain cases, these proteins appear to have re-converged on common solutions to carry out specific tasks, such as protein unfolding and degradation^2–4,14^. Our findings provide a clear example of this convergence by demonstrating how AAA+ quality control proteins from distantly related clades utilize distinct structural elements to power a conserved hand-over-hand mode of substrate conveyance.

## Materials and Methods

### Protein Expression and Purification

*E. coli* strain BL21 star (DE3) was used to express recombinant proteins. Cells were grown in LB broth (5 g yeast extract, 10 g tryptone, and 5 g NaCl per liter) supplemented with 50 µg/ml kanamycin. Cells were cultured at 37° C with shaking at 250 rpm. We have previously described the cloning and expression of *Y. pestis lon*, *hspQ*, and *y2853* genes^31^. The pET28b-*lon* plasmid was used to overexpress *Yersinia* Lon protein in *E. coli* strain BL21 star (DE3). Cells were cultured in LB containing 50 µg/ml kanamycin at 37° C to optical density at 600 nm (OD_600_) of 0.5. Protein expression was induced by 1 mM IPTG. Protein overexpression was carried out for 16 hours at 16° C. Cells were harvested by centrifugation at 3,700 xg and resuspended in buffer A (50 mM KHPO_4_ pH 7, 1mM EDTA, 1mM DTT, and 10% glycerol). After sonication, cleared cell lysate was prepared by centrifugation at 30,000 xg. Activated and buffer A equilibrated P11-cellulose resin was added to the cleared cell lysate to allow Lon binding. The column was washed with buffer A to remove unbound proteins, and bound Lon protein was eluted in 10 ml of elution buffer B (400 mM KHPO_4_ pH 7, 1mM EDTA, 1mM DTT, and 10% glycerol). Lon protein was further purified on a Source 15Q ion-exchange column using buffer C (50 mM Tris pH 7.5, 50 mM KCl, and 1 mM DTT). Bound Lon was eluted using a 20 column-volume linear-gradient (0-100%) of buffer D (50 mM Tris pH 7.5, 1 M KCl, and 1 mM DTT). Fractions containing Lon protease were pooled, concentrated, and loaded on S300 gel filtration column in buffer E (50 mM Tris pH 7.5, 100 mM KCl, 10 mM MgCl_2_, 1mM DTT, 20% glycerol). Aliquots of purified Lon were flash-frozen and stored at −80° C. The pET28b-*lon* plasmid served as a template for generating *lon* mutants using the quick-change site directed mutagenesis approach. Each Lon mutant protein was expressed and purified as described above.

HspQ and Y2853 were purified using a combination of Ni-NTA affinity, ion exchange, and size exclusion chromatography steps. BL21 star (DE3) harboring pET28b-hspQ plasmid was grown in LB containing 50 μg/ml kanamycin, and protein expression was induced with 1 mM IPTG at OD_600_ of 0.5. Cultures were allowed to grow for 3 h while shaking. Harvested cells were resuspended in lysis buffer (50 mM Tris pH 8, 1M NH_4_Cl, 2 mM beta-mercaptoethanol (β-ME), and 10 mM imidazole). After sonication, buffer-equilibrated Ni-NTA beads were added to cleared cell lysates. After 1 h end-to-end rocking at 4° C, unbound proteins were removed and the beads were washed extensively, and the bound proteins were eluted using a step elution with lysis buffer containing 250 mM imidazole. HspQ protein containing fractions were combined, buffer exchanged to buffer F (50 mM Tris pH 8, 50 mM KCl, 2 mM β-ME), loaded on a Source15Q column, and eluted using a 20 column-volume linear-gradient (0-100%) of buffer G (50 mM Tris pH 8, 1M KCl, 2 mM β-ME). Fractions containing HspQ were pooled, concentrated and loaded on a Superdex 75 column equilibrated in buffer H (50 mM Tris pH 8, 50 mM KCl, 2 mM β-ME). Protein aliquots containing 10% glycerol were flash-frozen and stored at −80° C.

### Proteolysis Assay

Each *in vitro* proteolysis assay reaction was carried out in Lon activity buffer (50 mM Tris-HCl pH 8, 100 mM KCl, 10 mM MgCl_2_, 1 mM DTT, and 10% Glycerol), ATP regeneration system (16 mM creatine phosphate, 0.32 mg/ml creatine kinase, and 4 mM ATP). Reactions contained 100 nM Lon hexamer (Lon_6_) and 10 *µ*M each. All reaction components except ATP regeneration system were mixed and incubated at 37° C. ATP regeneration system, pre-warmed at 37° C, was added to initiate the reaction. Aliquots at specific time were mixed with 2X-SDS sample buffer to terminate the reaction. Reaction products were resolved by electrophoresis on 15% Tris-Tricine gels and scanned using a Li-COR Odyssey scanner and quantified using the Image Studio software. Fraction of substrate remaining was estimated and the data was normalized to creatine kinase as loading control. Three biological repeats were performed for each Lon mutant and the data were fit to a straight line and the slope was extracted to calculate the rate of substrate degradation. GraphPad Prism software was used for data analysis. Mean and standard error of mean (SEM) was calculated by performing column statistics.

### *In vitro* ATP-Hydrolysis Assay

Coupled ATP hydrolysis assay was carried out in Lon activity buffer (50 mM Tris-HCl pH 8, 100 mM KCl, 10 mM MgCl_2_, 1 mM DTT, and 10% Glycerol). Reactions contained 100 nM Lon6, 1 mM NADH, 10 U/ml lactate dehydrogenase, 20 mM phosphoenol pyruvate, 10 U/ml pyruvate kinase, and 2 mM ATP. Lon and other reaction components were warmed separately at 30° C. Reactions were initiated by adding Lon, and NADH disappearance was monitored at 340 nm. Three biological repeats were performed for each Lon mutant and the indicated substrate protein and the data were fit to a straight line and the slope were extracted to calculate ATPase activation rates. GraphPad Prism software was used for data analysis. Mean and standard error of mean (SEM) was calculated by performing column statistics.

### Sample preparation for electron microscopy

LON^WB^ was diluted to a concentration of 0.95 mg/ml in 50 mM Tris pH 8, 75 mM KCl, 10 mM MgCl_2_, 1 mM TCEP, and 1 mM ATP. 4 µl of the sample were applied onto 300 mesh R1.2/1.3 UltrAuFoil Holey Gold Films (Quantifoil), that had been previously plasma treated for 30 seconds using a 15 mA current operating under atmospheric gases using a glow discharger (Electron Microscopy Sciences). Excess sample was blotted away for 4 s using Whatman No. 1 filter paper and vitrified by plunge freezing into a liquid ethane slurry cooled by liquid nitrogen using a manual plunger in a 4° C cold room whose humidity was raised to 95% using a humidifier.

### Electron microscopy data acquisition

Cryo-EM data were collected on a Thermo-Fisher Talos Arctica transmission electron microscope operating at 200 keV using parallel illumination conditions^38^. Micrographs were acquired using a Gatan K2 Summit direct electron detector, operated in electron counting mode applying a total dose of 51 e^-^/Å^2^ as a 44-frame dose-fractionated movie during a 11s exposure. The Leginon data collection software^39^ was used to collect 4071 micrographs at 36,000x nominal magnification (1.15 Å/pixel at the specimen level) with a nominal defocus range of −0.8 to −1.3 µm. Stage movement was used to target the center of four 1.2 *µ*m holes for focusing, and an image shift was used to acquire high magnification images in the center of each of the four targeted holes.

### Image processing

Real-time preprocessing was performed during cryo-EM data collection using the Appion processing environment^40^. Micrograph frames were aligned using MotionCor2^41^ and CTF parameters were estimated with CTFFind4^42^. Approximately 100,000 particles were selected from a subset of micrographs using a Difference of Gaussian (DoG)-based automated particle picker^43^. Particles were extracted using a box size of 128 pixels and the stack was binned by a factor of 2 for reference-free 2D alignment using an iterative multivariate statistical analysis with multi-reference alignment (MSA-MRA) in Appion. 2D classes representing orthogonal views of the Lon complex were used for template-based particle selection with FindEM^44^. A stack of 1,176,205 particles was created using a 256 pixel box size, which was scaled down by a factor of 4 using RELION 1.4^45^ for initial processing. An *ab initio* model was created in cryoSPARC^46^, low pass filtered to 60 Å and used as an initial model for 3D refinement of particles in RELION 2.0^47^. These particles refined to a reported resolution of 9.5 Å as estimated by Fourier Shell Correlation using a cutoff of 0.143, but the resolution was anisotropic, and the reconstruction exhibited artifacts from preferred orientation. The particles from this reconstruction were then sorted by classification without alignment into four classes, two of which, accounting for 65.9% of particles, displayed high-resolution features that did not contain anisotropic resolution artifacts. The 788,445 particles from these classes were merged and refined. The x & y shifts from this refinement were used to re-extract centered, unbinned particles using a box size of 256 pixels. The re-extracted particles were then refined and post-processed to produce a reconstruction with an estimated resolution of 3.7 Å at an FSC cutoff of 0.143. 2D Classification was used at this point to filter out false particle picks and noise, resulting in 749,413 particles that were further refined to an estimated overall resolution of 3.6 Å.

The cryo-EM density of the two seam subunits was poorly resolved in this reconstruction, so a soft-edged 3D mask was generated to encompass the step subunit for focused classification with three classes, resulting one class containing 10% of the particles that contained higher resolution step subunits and an overall reported resolution of 3.4 Å by FSC at 0.143. CTF refinement was performed to estimate per-particle defocus values using RELION 3.0^48^. The X and Y image shifts applied during data acquisition were used to group images for beam tilt estimation. Refining with local defocus and beam tilt estimation improved the overall reported resolution of our reconstruction to 3.0 Å by FSC at 0.143. Focused refinement of the masked step subunits and N-terminal domains improved the quality of the map in these regions to a reported resolution of 3.5 Å by FSC at 0.143. A final composite map of the focused and non-focused refinements was generated for atomic model building and refinement using the “vop max” operation in UCSF Chimera^49^.

### Atomic model building and refinement

A homology model was generated using the crystal structure of LON as a starting model using SWISS-MODEL^50^. This initial model was split into ATPase and protease domains and rigid body docked into the density of each of the subunits using UCSF Chimera^49^. Real-space refinement of the docked structures and *ab initio* model building were performed in COOT^51^. The seam ATPase subunits and flexible linker regions were modeled *ab initio* and an eight-amino acid polyalanine peptide, as well as ATP, ADP, and magnesium cofactor molecules were built into the density corresponding to substrate and nucleotide. Further refinement of the full hexameric atomic model was performed using real-space refinement in PHENIX^52^. This refined model served as a starting point to generate 100 models in Rosetta and the top five scoring models were selected for further refinement in Phenix and COOT using the multi-model pipeline^53^. UCSF Chimera and ChimeraX^54^ was used visualize the structure and to generate the figures. Only the top-scoring model is included in the figures.

## Acknowledgements

We thank J.C. Ducom at Scripps Research High Performance Computing and C. Bowman at Scripps Research for computational support, as well as B. Anderson at the Scripps Research Electron Microscopy Facility for microscopy support. We thank C. Sandate, A. Hernandez, M. Herzik, and M. Hirschi for assistance in atomic model building and refinement, as well as members of the Lander and Wiseman labs at Scripps Research for helpful discussions. M.S. is supported by the National Science Foundation Graduate Research Fellowship Program. G.C.L. is supported by the Pew Charitable Trusts as a Pew Scholar, an Amgen Young Investigator award, and by the National Institutes of Health (NIH) DP2EB020402. A.W.K. is supported by NIH AI-127533. R.L.W. is supported by the NIH NS095892. G.C.L. and R.L.W. are supported by NIH AG061697. Computational analyses of EM data were performed using shared instrumentation funded by NIH S10OD021634 to G.C.L.

## Data Availability

The EM density map and the top-scoring atomic model obtained from the multi-model pipeline^38^ have been deposited at the Electron Microscopy Data Bank and Protein Data Bank under accession numbers EMDB: 20133 and PDB: 6ON2.

## Supplementary Materials

**Figure S1.**
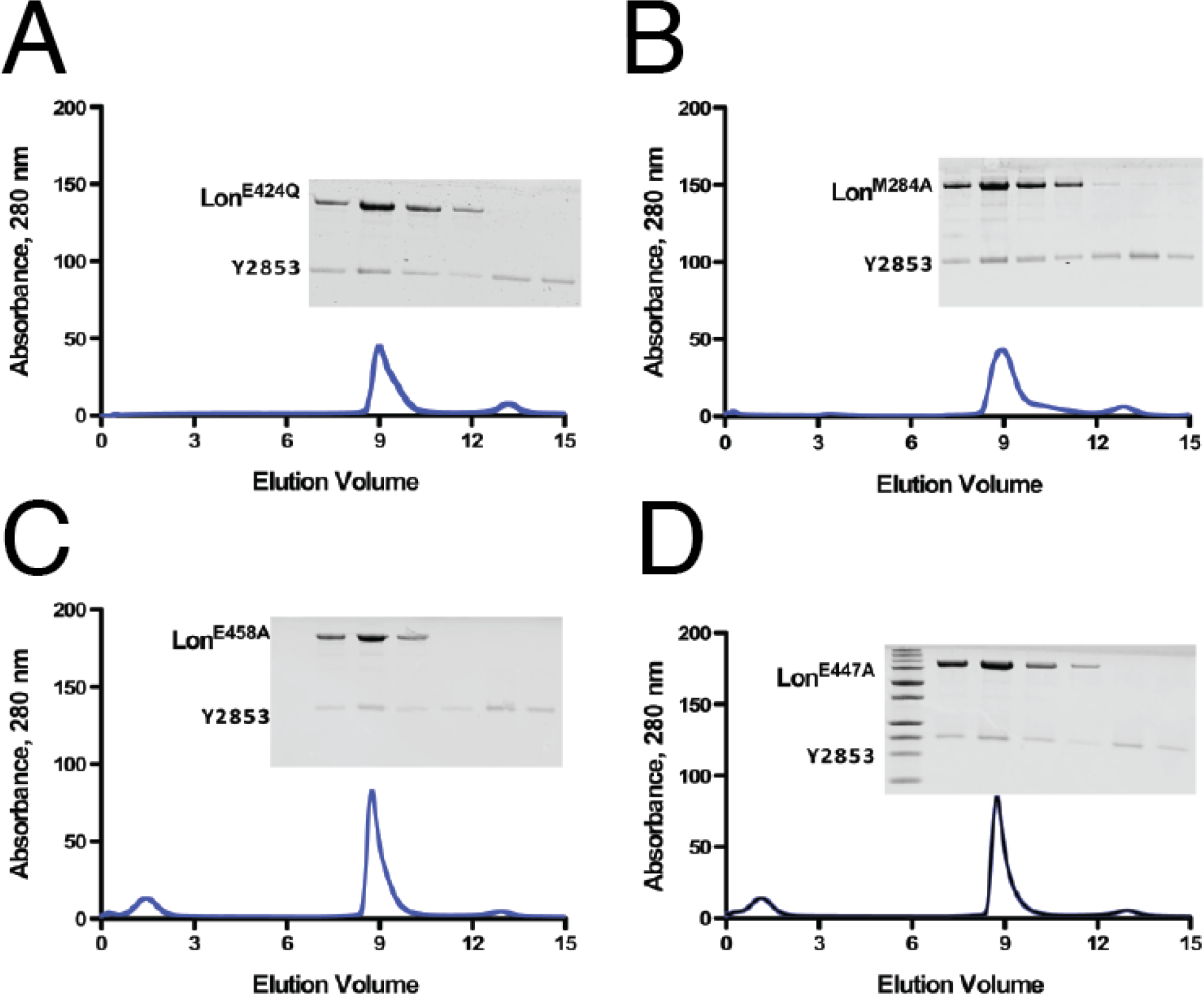
Purification of substrate-bound Lon complexes. Size-exclusion chromatography (SEC) traces showing Y2853 substrate-bound Lon complexes used for structural and biochemical analyses eluting around 9 mL for **A.** WT Lon bearing a E424Q (Walker B) mutation used for structural studies, **B.** M284A mutation, **C.** E458A mutation, and **D.** E447A mutation, indicating a complex size of 90 kDa. Overlaid onto traces are SDS-PAGE stained with Coomassie Brilliant Blue showing denatured contents of the elution fractions from the SEC experiment.

**Figure S2.**
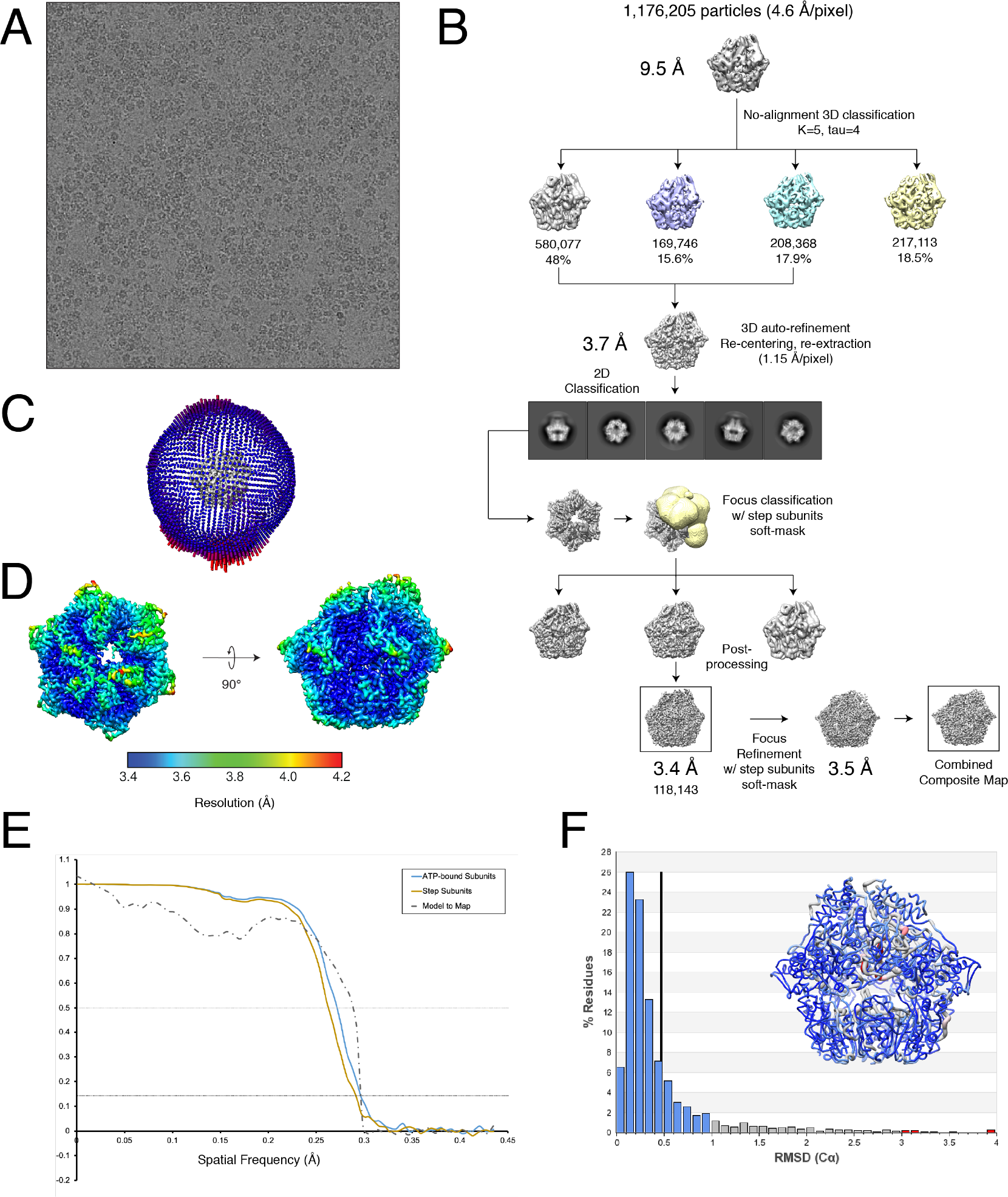
Structure determination workflow and validation. **A.** Representative micrograph from cryo-EM data collection. **B.** Cryo-EM data processing scheme followed using RELION 2.1b software to obtain the final 3D reconstruction of substrate-bound Lon. Final steps included a focused refinement of the final reconstruction using a soft-mask over the step subunits and stitching together the focused region with the remainder of the final map. The combined composite map was used for atomic model building and refinement. **C.** Euler angle distribution plot of the 118,143 particles used in the final reconstruction and combined composite map. **D.** Final reconstruction filtered by local resolution calculated using BSOFT^56^. The final EM density carries a range of resolutions, from 2.8 Å at the core of the complex to > 4.2 Å in more flexible regions such as the step subunits. **E.** Fourier Shell Correlation (FSC) of the final reconstruction (blue solid line) used to compose four ATP-bound subunits in the combined composite map, the focused refinement map (solid brown line) used to compose two seam subunits in the combined composite map, and the top-refined atomic model vs. the composite combined map (dotted black). **F.** A histogram showing the per-residue Cα RMSD values calculated from the top ten refined atomic models using the multi-model pipeline^38^. A vertical black bar represents the mean per-residue Cα RMSD value, and a worm representation of Lon colored according to the per-residue Cα RMSD values (in Å) is overlaid onto the histogram.

**Figure S3.**
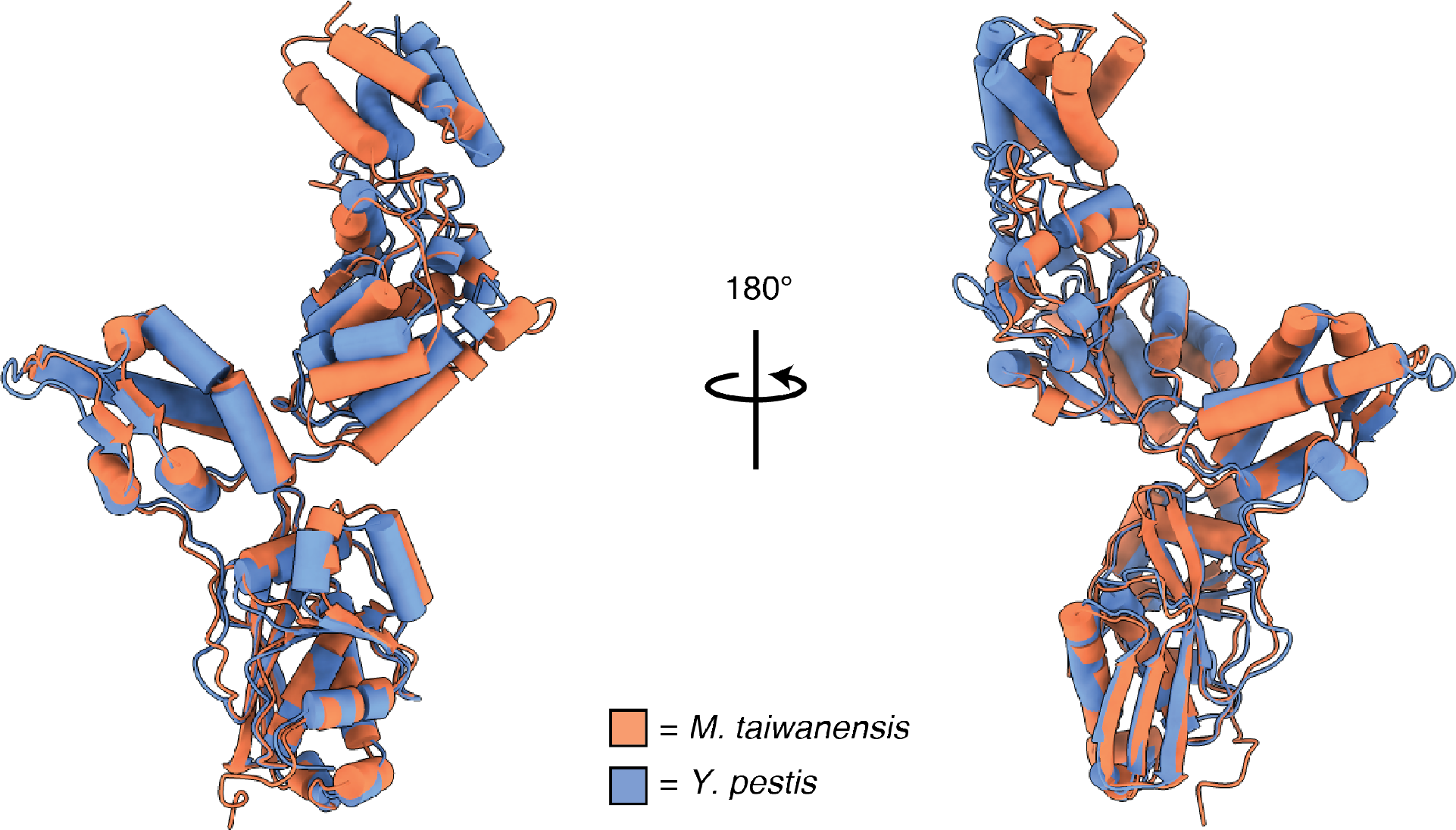
Comparison to prior x-ray structure. Secondary structural alignment of the ATP2 subunit showing strong resemblance to the nucleotide-free protomer structure of a previously determined crystal structure of substrate-free *M. taiwanensis* Lon solved using x-ray crystallography (PDB:4YPL). The average Cα RMSD values showed minimal deviations in protease and NTD^3H^ of the two structures (0.777 and 1.011 Å, respectively) whereas the ATPase domains are slightly more variable (1.256 Å). These results show the similarity of the two protomers and consistency of the secondary and tertiary structures of an individual subunit. Higher Cα RMSD values for the ATPase domains are likely due to differences in conformations between a nucleotide and substrate-bound to a nucleotide and substrate-free structure.

**Figure S4.**
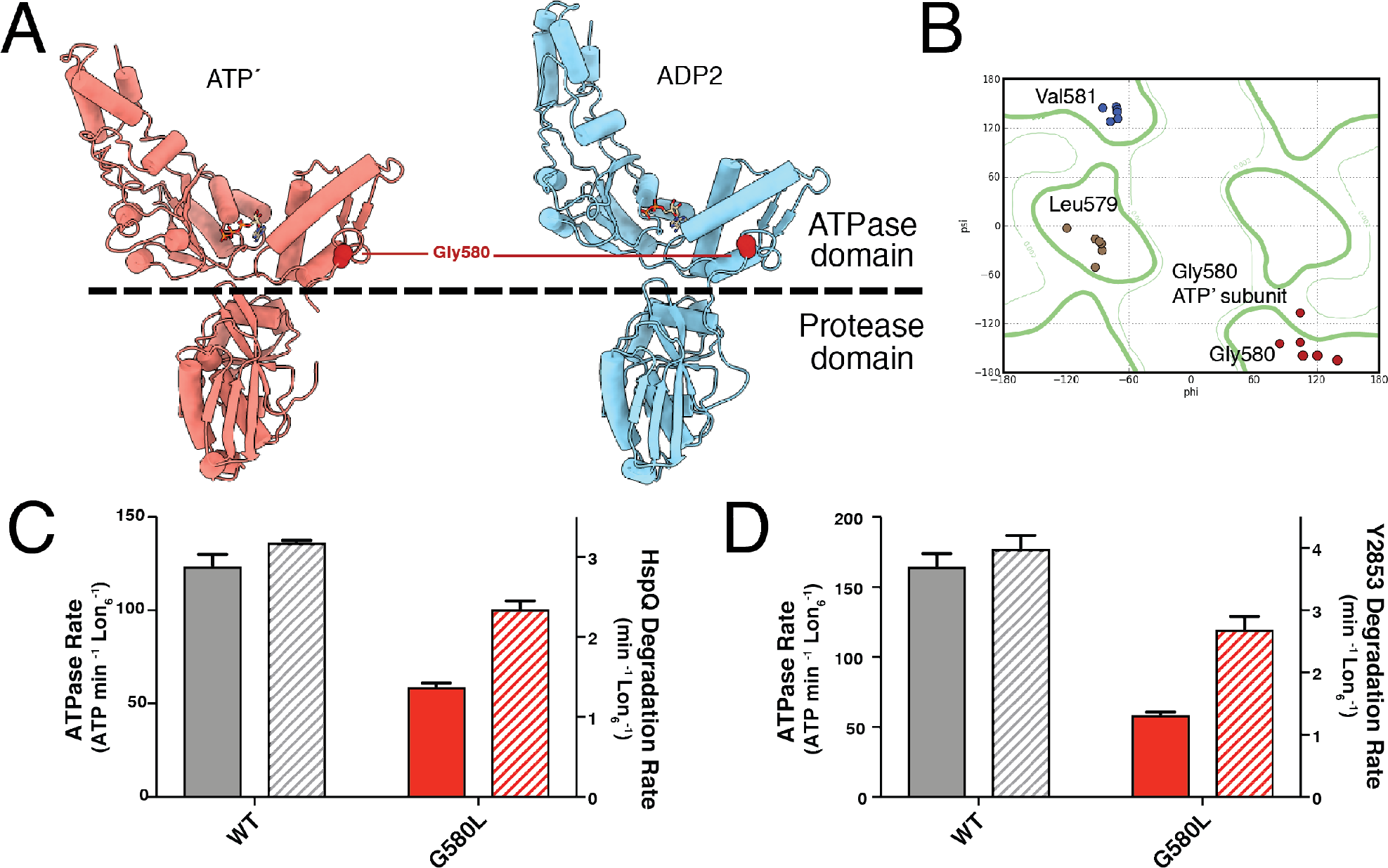
A flexible glycine-containing linker allows for large-scale conformational changes of ATPase domains. **A.** Side-by-side comparison of ATP′ and ADP2 subunits in the same orientation shows rigid body movements of the ATPase domain between the two subunits while the protease domains remain stable. A glycine residue (G580) in the flexible interdomain linker is highlighted in red. **B.** Ramachandran plot of Gly580 and surrounding residues shows a dihedral angle transition between ATP′ and ADP2 subunits, indicating a hinge-like role of the glycine residue. **C-D.** Effects of mutating Gly580 to leucine on substrate-induced ATPase activity and degradation of known Lon substrates, HspQ and Y2853. G580L mutant causes defects in both substrate-induced ATPase and substrate degradation rates. These results indicate that hinge-like movements in this residue are critical for both hydrolysis and translocation in Lon, whilst only mildly affecting ATPase rates in other AAA+ proteases. These differences between AAA+ proteases are likely due to the unique allosteric mechanism present in Lon.

**Figure S5.**
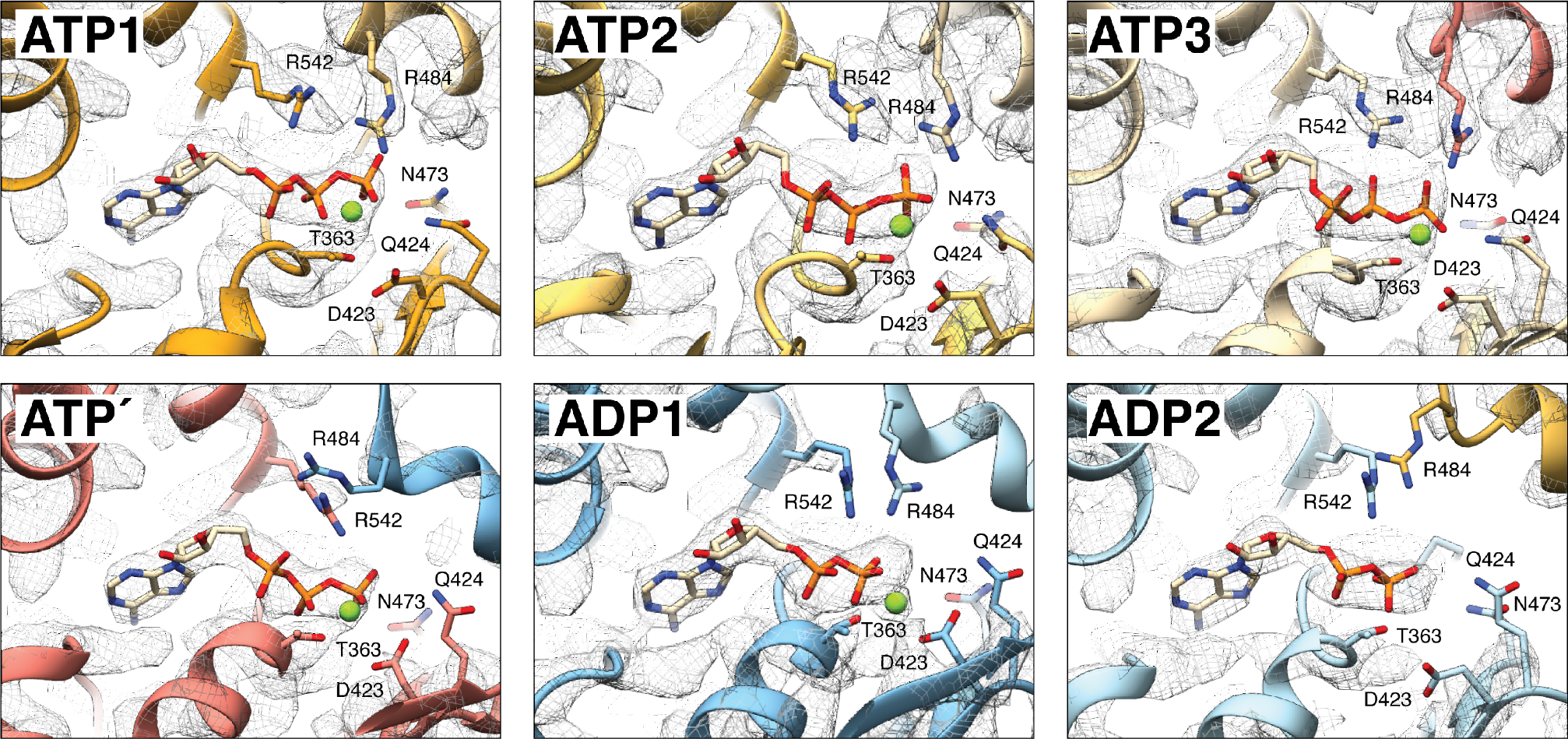
Distinct nucleotide densities in the nucleotide-binding pocket. Detailed views of the nucleotide binding pockets of all six subunits showing the quality of the cryo-EM density in this region. cryo-EM density of each subunit is shown using an isosurface mesh representation contoured at sigma = 3.3. The quality of the EM density enables unambiguous assignment of nucleotide state in each of the subunits. ATP1, ATP2, ATP3, and ATP′ subunits possess strong density for nucleotide corresponding to ATP with gamma phosphate coordination by a magnesium cofactor. In contrast, the nucleotide density in the ADP1 and ADP2 subunits corresponded to ADP molecules, as there was no apparent density for gamma phosphates nor magnesium cofactors. These data suggest there are two coexisting nucleotide states within the substrate-bound structure with four sequential subunits bound to ATP and two seam subunits bound to ADP.

**Figure S6.**
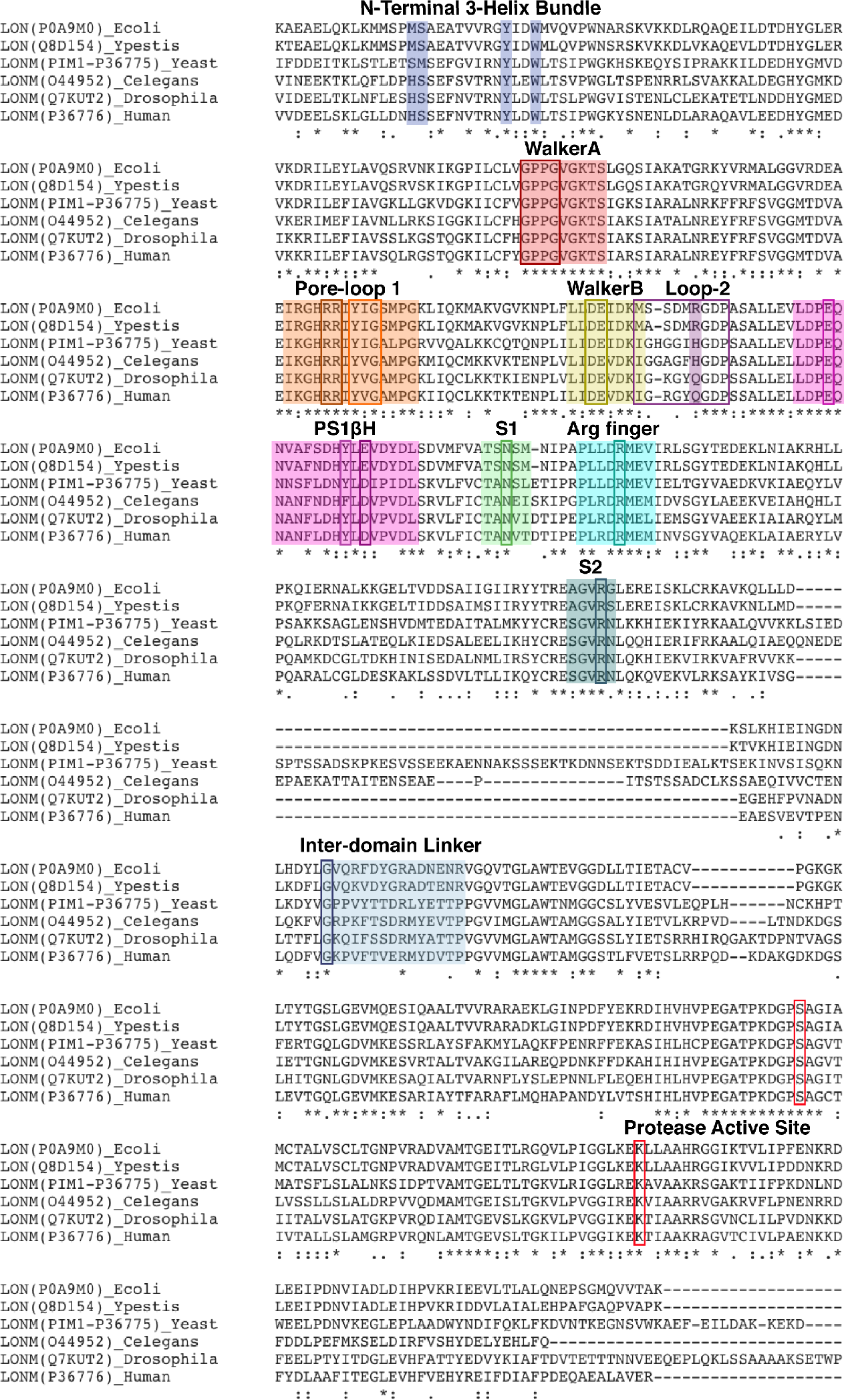
Conservation of the allosteric mechanism across Lon proteases. Clustal W alignment of the Uniprot sequences of the NTD^3H^ and ATPase domains of Lon homologs, including cytoplasmic *E. coli* and *Y. pestis* Lon and mitochondrial Lon from yeast (Pim1), *C. elegans*, *Drosophila melanogaster*, and humans. This alignment suggests strict mechanistic conservation amongst Lon proteins in diverse model organisms and environments, as we found all key residues identified in our structure to be strictly conserved in all sequences studied: the N-terminal 3-Helix Bundle (M294, S285, Y294, and W297 in light blue), P-loop (purple box), pore-loop 1 aromatic-hydrophobic residues (pink box), coordinating acidic residues in the Walker B motif (dark blue box), pre-sensor 1 beta hairpin insertion with conserved residues E447, Y456, and E458 in boxes, sensor-1, a trans-acting arginine finger (maroon box), as well as a cis-acting arginine finger present in sensor-2 (green).

**Figure S7.**
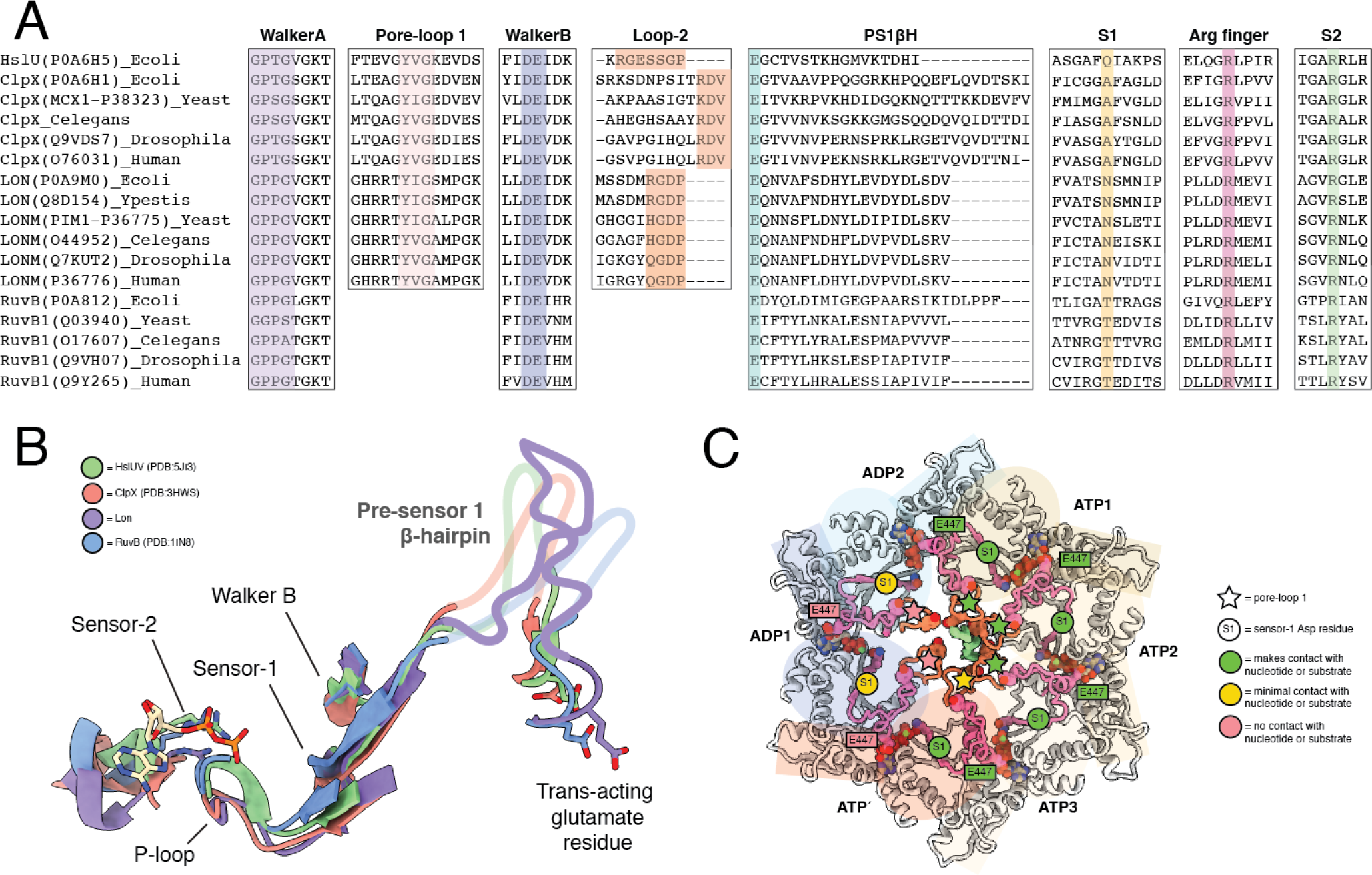
Conservation of allosteric mechanism across HCLR clade proteins. **A.** Alignment of conserved regions in the AAA+ domains of HCLR clade proteins, HslU, ClpX, Lon, and RuvB. This alignment suggests mechanistic conservation amongst HCLR clade proteins, as we found all key residues identified in our structure to be strictly conserved in all sequences studied: P-loop (purple in Walker A motif), pore-loop 1 aromatic-hydrophobic residues in protein translocases (pink), coordinating acidic residues in the Walker B motif (blue), a pre-sensor 1 beta hairpin insertion with a trans-acting glutamate residue (teal) at its N-terminus, sensor-1, a trans-acting arginine finger (maroon), as well as a cis-acting arginine finger in sensor-2 (green). **B.** Structural conservation of essential elements of the proposed allosteric mechanism amongst HCLR clade proteins, including HslU (PDB:5JI3), ClpX (PDB:3HWS), Lon (PDB:6ON2), and RuvB (PDB:1IN8). **C.** A top view of the substrate-bound Lon with each ATPase domain highlighted by an ellipse and rectangle on the large and small subdomains, respectively. Key structural elements are highlighted in hot pink: residues that comprise the pore-loop 1, PS1βH, sensor-1, and a trans-acting glutamate residue. The mechanism of allostery in Lon involving these shared elements is likely conserved across all HCLR clade proteins.

**Table S1.** Cryo-EM data collection, refinement, and validation statistics.

| Y. pestis Lon protease, PDB ID: 6ON2, EMDB ID: 20133 |  |  |
| --- | --- | --- |
| Data Collection |  |  |
| Microscope | Talos Arctica |  |
| Camera | K2 Summit |  |
| Magnification (nominal) | 36,000 |  |
| Magnification (at detector) | 43,478 |  |
| Voltage (kV) | 200 |  |
| Data acquisition software | Leginon<sup>39</sup> |  |
| Exposure navigation | image shift |  |
| Total electron exposure (e<sup>-</sup>/Å<sup>2</sup>) | 52 |  |
| Exposure rate (e<sup>-</sup>/pixel/sec) | 5.3 |  |
| Number of frames | 44 |  |
| Defocus range (μm) | 0.8 to 1.8 |  |
| Pixel size (Å/pixel)* | 1.15 |  |
| Micrographs collected (no.) | 4,071 |  |
| Micrographs used (no.) | 4,071 |  |
| Total extracted picks (no.) | 1,176,206 |  |
| Refined particles (no.) | 1,176,206 |  |
| Reconstruction |  |  |
| Final Particles (no.) | 118,143 |  |
| Estimated accuracy of rotations/translations | 0.67/0.32 |  |
| Symmetry | C1 |  |
| Resolution (global) (Å) |  |  |
| FSC 0.5 (unmasked/masked) | 4.2/3.3 | ADP1-ADP2: 4.3/3.8 |
| FSC 0.143 (unmasked/masked) | 3.5/3.0 | ADP1-ADP2: 3.8/3.5 |
| Local resolution range (Å) | 2.8 - 4.2 |  |
| Map sharpening B-factor (Å<sup>2</sup>) | -51 | ADP1-ADP2: -105 |
| Model composition |  |  |
| Protein residues | 3,146 |  |
| Ligands | 6 |  |
| Refinement* |  |  |
| Refinement package | Phenix<sup>52</sup> |  |
| MapCC (global/local) | 0.79/0.70 |  |
| B-factors (Å<sup>2</sup>) |  |  |
| Protein residues | 54.4 |  |
| Ligands | 37.2 |  |
| R.m.s. deviations |  |  |
| Bond lengths (Å) | 0.01 |  |
| Bond angles (°) | 0.83 |  |
| Validation |  |  |
| MolProbity score | 1.63 |  |
| Clashscore | 6.69 |  |
| Poor rotamers (%) | 0.38 |  |
| C-beta deviations | 0 |  |
| Ramachandran plot |  |  |
| Favored (%) | 96.14 |  |
| Allowed (%) | 3.83 |  |
| Outliers (%) | 0.03 |  |
| Mean per-residue Cα RMSD (Å) | 0.46 |  |
| Per-residue Cα RMSD range (Å) | 0.3-5.32 |  |
| CaBLAM outliers (%) | 0.02 |  |
| EMRinger<sup>55</sup> score | 3.0 |  |

\*Atomic statistics and validation provided for best-scoring atomic model

